## Supplementary table 1 and S1 extended methods; S2 follow up studies for "Genetic differentiation at extreme latitudes in the socially plastic sweat bee *Halictus rubicundus*"

### Supplementary information

**Table 1:** The three sets of primers that were used.

| Set | Primer name | Range | Dye colour | GenBank accession number | Primer sequences (5'-3') (F primers are labelled on 5' end) |
|---|---|---|---|---|---|
| S1 | rub02 | 168-190 | Blue | BV729095 | F: CCAGCCGGCCAACGTTGC<br>R: CGGAGCTGAAAACTCAATTACAG |
| S1 | rub30 | 150-200 | Yellow | BV729098 | F: GATCCGCTTTCAACCGTCCG<br>R: GTGAGCTGGGTCCGGCGAG |
| S1 | rub35 | 177-273 | Red | BV729108 | F: GATGACGCAGTACGAACGG<br>R: GCTTCGACGTATGATTATCC |
| S1 | rub37b | 88-94 | Blue | BV729099 | F: GATTTTTCTCGCGTACCTCTGC<br>R: CACTGCGATTCCGGGTTGTCC |
| S1 | rub59 | 185-211 | Green | BV729101 | F: GTGACCAGGTGCGCTCGTTAC<br>R: CCGTGTCCCCAGCTCCGTTTC |
| S2 | rub04 | 197-205 | Red | **?** | **?** |
| S2 | rub06 | 238-284 | Blue | BV729097 | F: GTCTGGCGGAAGTCTACGTGC<br>R: CAAGTTCGGTGCGTTAGATAATG |
| S2 | rub55 | 133-155 | Yellow | BV729100 | F: GCTATAAAAGGCGAAACGGGTG<br>R: CTCCTATCCGGTTGACATTGCC |
| S2 | rub60 | 123-149 | Blue | BV729102 | F: GCAAACACACCGCTAATGACATG<br>R: GCCGACAGGTTTGCAGCATGAG |
| S3 | rub73 | 165-195 | Yellow | **?** | **?** |
| S3 | rub80 | 100-180 | Green | BV729107 | F: CCGGTCGGAGGTGTGTTC<br>R: CATACTTCCTTCCTAGCATTCG |
| S3 | rub61 | 180-202 | Red | BV729103 | F: GACGCGGAAATAGAAAAGTTG<br>R: CTAATGCATCGGGCCAACTG |
| S3 | rub72 | 149-177 | Blue | BV729104 | F: GCATTTATTCCGTCGCGACTC<br>R: CGGTGGCGGGGCTCGTAATG |

S1 Extended methods

**Peak scoring**

In this research project the values of each peak within this Geneious file were scored in Geneious v. R11 (https://www.geneious.com) in a marker-focused manner (i.e. looking at a single microsatellite marker for all individuals before moving on to the next marker). Scored values represent the number of microsatellite repeats for each marker, different values for the same microsatellite marker represent different alleles. Peaks were scored for quality and some individuals were scored multiple times to ensure consistency. Some individuals were genotyped a second time if initial PCRs partially or fully failed to amplify DNA or if reliable scores could not be determined due to stutter around peaks or other common issues with repeatability

While scoring the peaks it was important to determine which of the peaks represents the allele and which are noise. A peak needs to meet several requirements to be scored as the allele peak. It should be distinct from the background. In the case of heterozygosity, the second peak should be lower than the first peak. Smaller peaks lead up to the final peak. These leading peaks are a good inclination that the final peak of that series represents the allele (figure S1). Small peaks in front of the allele peak can be explained due to adenylation.

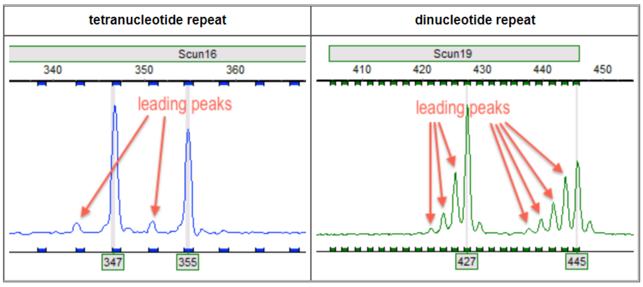


**Figure S1:** What leading peaks look like in Geneious [22].

As these microsatellites were created over a decade before this experiment was performed, it is to be expected that some of the peak ranges have shifted. This was true with almost all locus ranges, which were thus altered if a lot of obvious peaks were found out of range in multiple individuals. The S1 blue locus range contained two markers in the same colour. Alteration of these locus ranges were handled with extra caution, as I could have accidentally increased the range in such a way that an allele from the first marker would have been scored in the second one.

**GeneAlEx analyses**

Nei's unbiased genetic distance may give spurious results when homozygosity and sample size is small, so it was decided to use the normal Nei's genetic distance.

Several analyses were performed via GeneAlEx, which were calculated in the following way.

$F_{ST}$ is the a measure of population differentiation due to genetic structure. $F_{ST} = (H_E - H_O)/H_E$.

$H_O$ is the observed heterozygosity. $H_O$ = Number of heterozygotes / Total number of individuals

$H_E$ is the expected heterozygosity. $H_E = 1 - \text{Sum } pi^2$.

Where pi is the frequency of the $i^{th}$ allele for the population and Sum $pi^2$ is the sum of the squared population allele frequencies.

Nei's genetic distance is another measure similar to Fst. Both are true in our case, so this wasn't used.

$$Nei\ I = \frac{J_{xy}}{\sqrt{(J_x J_y)}}$$

Where, $J_{xy} = \sum_{i=1}^{k} p_{ix} p_{iy}$, $J_x = \sum_{i=1}^{k} p_{ix}^2$, and $J_y = \sum_{i=1}^{k} p_{iy}^2$.

Where *I* is Nei's Genetic Identity, and $p_{ix}$ and $p_{iy}$ are the frequencies of the *i*-th allele in populations *x* and *y*. For multiple loci, $J_{xy}$, $J_x$ and $J_y$ are calculated by summing over all loci and alleles and dividing by the number of loci. These average values are then used to calculate *I*.

*Nei D* = -ln(*I*). Gives the Nei genetic distance value.

S2 Follow-up studies

I suggest a follow-up research project. By placing marked bees from a different environment, such as Cornwall, in this Scottish environment and sampling the percentage of marked bees at the start and at the end of the growing season, one could compare if there is a difference in fitness between these bees, as the percentage of marked bees would decrease if they have a lower fitness and thus lower survival rate compared to the bees that are native to this environment. If the percentage is the same, another study could be done. In this study, one could sequence the genome of some Migdale bees, and some bees from a different location and then put these bees originating from another environment in the Migdale environment. By then sequencing the next generation of bees a year later, one could check if any breeding took place between the bees. If the bees can survive in Migdale, but have a phenological barrier, then is expected to not find a lot of breeding between the two types of bees. If they do survive and breed, then neither phenological differences or differences due to local adaptation are expected to cause the inhibition of gene flow. It could then simply be that the genetic distance is due to a physical barrier in the form of Scottish highlands. One could of course also place the Migdale bees in a southern environment and then run similar tests.

Another possible way to test precisely where this barrier occurs, is by genotyping populations from all latitudes throughout the UK, and comparing them. However this is very time-consuming and expensive. The other proposal is more realistic, and could also explain what the barrier to gene flow is.
